# Contrasting Phyllosphere Mycobiome in two Lycopodiaceae Plant Species: Unraveling Potential HupA-Producing Fungi and Fungal Interactions

**DOI:** 10.1101/2024.01.18.576228

**Authors:** Liqun Lin, Cheng Li, Chiung-Chih Chang, Ran Du, Jiaojiao Ji, Li-Yaung Kuo, Li Wang, Ko-Hsuan Chen

## Abstract

**Background:** Huperzine A (HupA) is a natural lycopodium alkaloid renowned for its efficacy in treating neurodegenerative diseases such as Alzheimer’s disease. It specifically occurs in the Huperzioideae subfamily of Lycopodiaceae. Fungi associated with Huperzioideae species are potential contributors to HupA biosynthesis, offering promising prospects for HupA production. However, limited knowledge of fungal diversity in lycophytes, coupled with decreased HupA production over time of fungal strains, has impeded the discovery and applications of HupA-producing fungi. Here, we investigated huperzine concentrations and the mycobiome across various tissues of two Lycopodiaceae species, *Huperzia asiatica* (a HupA producer) and *Diphasiastrum complanatum* (a non-HupA producer). Our objectives across the tissues of the two species are to unveil the distribution of potential HupA-producing fungi and elucidate fungal interactions within the mycobiome, aiming to uncover the role of HupA-producing fungi and pinpoint their potential fungal facilitators.

**Results:** Among the tissues, *H. asiatica* exhibited the highest HupA concentration in apical shoots (360.27 μg/ml) whereas *D. complanatum* showed no HupA presence in any tissue. We obtained 441 Amplicon Sequence Variants (ASVs) from *H. asiatica* and 497 ASVs from *D. complanatum*. The fungal communities in bulbils and apical shoots of *H. asiatica* were low in diversity and dominated by Sordariomycetes, a fungal class harboring the majority of reported HupA-producing fungi. Integrating bioinformatics with published experimental reports, we identified 27 potential HupA-producing fungal ASVs, primarily in *H. asiatica*, with 12 ASVs identified as hubs in the fungal interaction network, underscoring their pivotal roles in mycobiome stability. Members of certain fungal genera, such as *Penicillium*, *Trichoderma*, *Dioszegia*, *Exobasidium*, *Lycoperdon* and *Cladosporium* exhibited strong connections with the potential HupA producers in *H. asiatica*’s network rather than in *D. complanatum*’s, implying their prospects as fungal facilitators in enhancing HupA production.

**Conclusions:** This study advances our knowledge of fungal diversity in Lycopodiaceae and provides insights into the search for potential HupA-producing fungi and fungal facilitators. It highlights the importance of exploring young tissues, and emphasizes the ecological interactions that promote the fungi-mediated production of complex bioactive compounds, offering new directions for research in fungal ecology and secondary metabolite production.

## Background

The phyllosphere mycobiome refers to fungal communities that inhabit both the surface (epiphytes) and interior (endophytes) of above-ground plant tissues [1]. While some phyllosphere fungi are known for their ability to promote plant growth and health, the functions of most of them remain understudied [2]. Recently, endophytic fungi have garnered attention as potential sources of bioactive compounds, especially in medicinal plants that harbor a wide range of endophytes [3–5]. Research has shown that some endophytic fungi can exert beneficial effects on host metabolism by producing, inducing, or modifying plant-derived natural products [6]. Biosynthesis of several well-known plant-derived bioactive compounds facilitated by endophytic fungi include taxol, camptothecin, podophyllotoxin and vincristine [7, 8]. These findings provide a promising avenue for developing an environmentally friendly, relatively simple, and cost-effective alternative method for producing bioactive compounds [3, 9, 10].

Huperzine A (HupA) is a natural lycopodium alkaloid isolated from the traditional Chinese medicine (TCM) “QianCengTa” [11]. This compound has received significant attention for its effect in the treatment of Alzheimer’s disease, the leading cause of dementia in older adults, which afflicts millions worldwide [12]. HupA has also shown promise in other neurodegenerative diseases and aging-related treatments [13, 14]. The increase of ageing populations is leading to an escalating demand for HupA, resulting in a supply-demand imbalance. Currently, HupA is primarily obtained through natural extraction and chemical synthesis. The Huperzioideae subfamily, belonging to the Lycopodiaceae family, is the only known natural source of HupA, whereas its sister lineages, Lycopodioideae and Lycopodielloideae (Supplementary Fig. S1), do not produce HupA [15, 16]. Extracting HupA from Huperzioideae species faces several challenges, including limited abundance, long growth cycle, low HupA content, and difficulties in cultivation [17]. Moreover, environmental deterioration and overexploitation of wild *Huperzia serrata* (the plant source of TCM “QianCengTa”) populations have raised apprehensions about the sustainability of HupA supply [18]. Chemical synthesis of HupA is also limited due to its complex structure, low productivity, and the fact that natural HupA is much more effective than its synthetic analogs [19].

Recent research has highlighted the potential of endophytic fungi in HupA production, offering a promising alternative to the challenges associated with natural extraction and chemical synthesis. The mycobiome of Huperzioideae species may play a key role in HupA production, with some fungal species potentially inducing HupA accumulation in the host plants [20–22]. Various endophytic fungi have been found to produce HupA independently in culture, without the need for cooperation with the host plant (Supplementary Table S1). These fungi offer a sustainable avenue of HupA production without large-scale cultivation or harvesting of the host plant. Their growth and metabolism can also be manipulated to increase the yield of HupA, making these fungi commercially valuable [23].

However, as of yet, none of the identified HupA-producing fungi have successfully been employed for HupA industrial production. Despite of the significant ecological values of Lycopodiaceae in various ecosystems [24], research on the Lycopodiaceae mycobiome has primarily been focused on a few well-known HupA-producing species, resulting in a limited understanding of fungal diversity within the broader range of Lycopodiaceae species, thus impeding the discovery of new fungal species with high HupA production capacity [21, 25]. Another major hurdle is the repeated sub-culturing of endophytic fungi leads to diminished or complete shut-down of HupA production over generations [26, 27], which could partially be attributed to the absence of other interactive fungi inhabiting the same ecological niche. The current process of identifying potential fungi producing specific secondary metabolites revolves around isolation, purification, and subculturing, often disregarding the probable impact of endophytic fungal interactions on secondary metabolite production [28]. Additionally, most strains isolated from the host plant are discarded without further exploration [29]. Nevertheless, several studies have demonstrated that co-cultivation of different fungi sharing the same ecological niche can significantly enhance the production of secondary metabolites in target fungi [30, 31]. However, co-cultivation applications for the production of fungal HupA have not been reported so far. Thus, investigating the interactions among fungi associated with Huperzioideae species, particularly those that produce HupA, could provide valuable insights into the correlation between metabolite content and fungal interaction networks.

In this study, a high-throughput sequencing method was employed to investigate the diversity and composition of endophytic fungal communities in various tissue parts of two lycophytes, *Huperzia asiatica* (producing HupA) and *Diphasiastrum complanatum* (not producing HupA). We utilized two approaches, sequence clustering and correlation analysis, to identify the potential HupA-producing fungi in *H. asiatica*. Furthermore, we employed networks analysis to reveal fungal interactions within the mycobiome. Overall, our goal is to advance our understanding of the intricate interactions between phyllosphere fungi and natural product biosynthesis, and thus to promote the application of fungi in the industrial production of plant-derived bioactive compounds.

## Methods

### Plant materials

The HupA-producing plants *H. asiatica* were collected in September 2019 from Changbai Mountain National Nature Reserve in Antu County, Jilin Province, China (41°41’49“-42°25’18”N, 127°42’55“-128°16’48“E, 2000-3000 m above sea level). The non-HupA producing plants *D. complanatum* were collected in June 2020 from Taipingshan National Forest Recreation Area in Yilan County (24° 30’ 10“ N, 121° 31’ 55“ E, 2000 m above sea level). Six tissue types of *H. asiatica* were sampled, including apical shoots, bulbils, young leaves, normal leaves, stems, and sporangia (Fig. 1A). Normal leaves and stems of *D. complanatum* were sampled (Supplementary Fig. S2). Three plant individuals were collected for the following analyses.

**Fig. 1.**
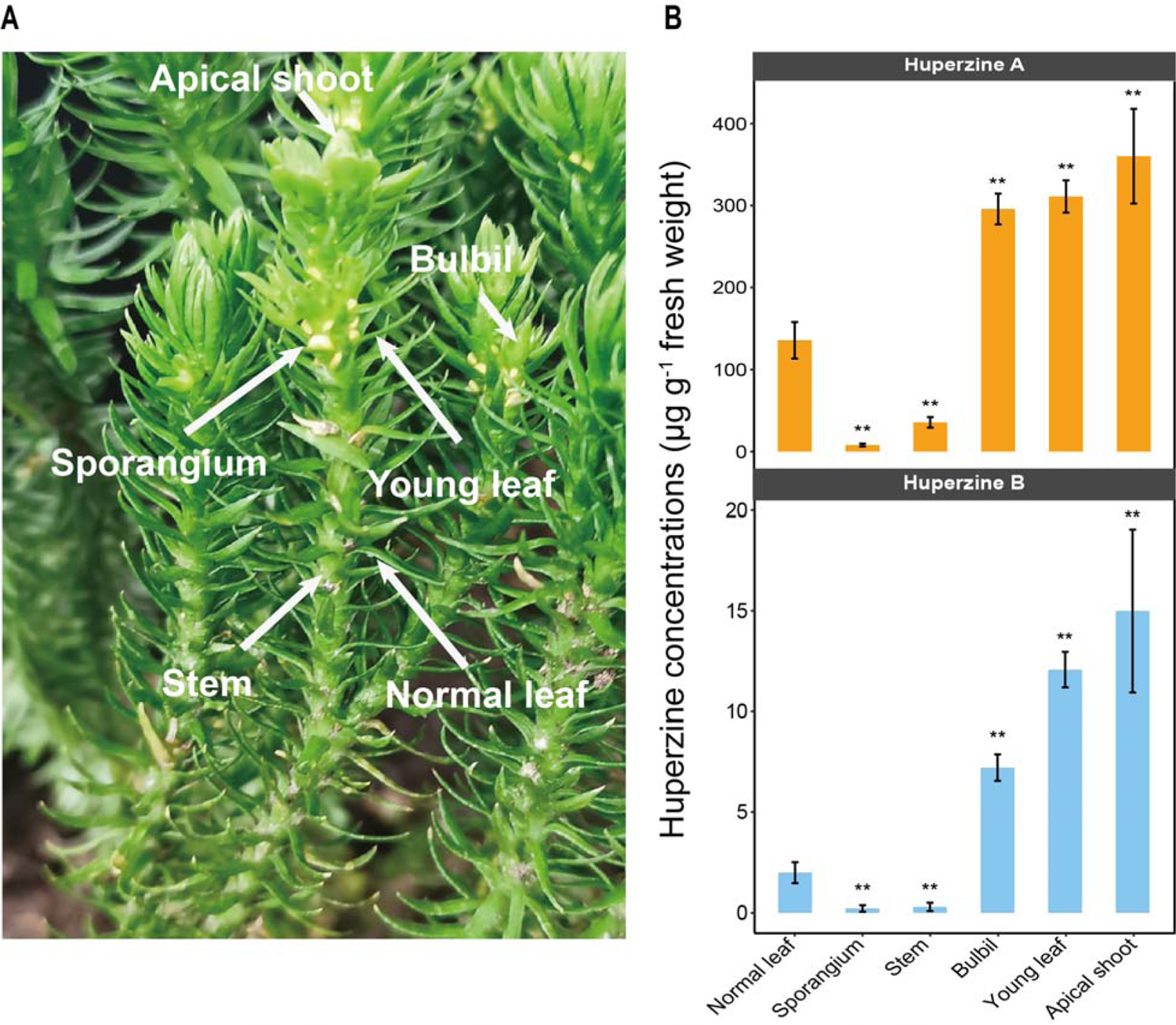
Huperzine concentrations. **(A)** Tissues of *Huperzia asiatica* plants (apical shoot, bulbil, young leaf, normal leaf, stem and sporangium) are used to quantify huperzine contents and to extract DNA for subsequent amplicon sequencing analysis. **(B)** The huperzine concentrations (microgram per gram fresh weight) in diverse tissues (Wilcoxon signed rank test, ** = *P* < 0.01). Initial concentration data are provided in Supplemental Table S2.

### LC-MS detection of Huperzines

All common chemicals and reagents were obtained from commercial vendors. Huperzine standards were purchased from commercial vendors: Huperzine A (ApexBio Technology LLC, Houston, TX, USA) and Huperzine B (MilliporeSigma, Burlington, MA, USA).

To measure the huperzine concentrations in plant tissues, samples were ground to fine powder in liquid nitrogen, lyophilized to dryness, and weighed to calculate the dry mass. Prepared samples were analyzed with an AB Sciex SCIEX TripleTOF 6600 plus system (AB SCIEX, MA, USA). 2 µL of extracts were injected and separated by an AB Sciex ExionLC system (AB SCIEX, MA, USA) equipped with an ACQUITY UPLC™ BEH C18 column (Waters Ltd., MA, USA) (150 × 2.1 mm, 2.6 μm). Water with 0.1% formic acid was used as mobile phase A, and ACN with 0.1% formic acid was used as mobile phase B, and the flow rate was 0.2 mL/min. A 20 min gradient method was used with mobile phase B gradually increased from 3% to 97%. Electrospray ionization (ESI) positive mode was used for detection, and the source conditions were set as follows: Ion Source Gas1 (Gas1) of 60, Ion Source Gas2 (Gas2) of 60, curtain gas (CUR) of 30, source temperature of 600°C, and IonSpray Voltage Floating (ISVF) 5500 V. Data with mass ranges of m/z 120-1800 was acquired under data dependent MSMS acquisition mode. In auto MS/MS acquisition, the instrument was set to acquire over the m/z range 25-1000 Da. The product ion scan parameters were set as follows: the collision energy (CE) was fixed at 35 V with ± 15 eV, the declustering potential (DP) was 60 V (+). Data was then analyzed by a SCIEX OS software (version 1.7, AB SCIEX, MA, USA). The presence of HupA and B in these samples was confirmed via both retention time and MS/MS data, and the concentration was calculated by calibration curves made with standards. Huperzine concentrations in *H. asiatica* across different tissues are shown in Supplementary Table S2.

### Amplicon sequencing

After collection, the samples were stored in silica gels until DNA extraction. We used a different number of pieces per DNA extraction for different tissue types due to the differences in their sizes (approximately one 2.5 cm stem of *D. complanatum* and *H. asiatica*, one 2.5 cm leaf of *D. complanatum*, five 0.5 cm bulbil/apical shoot of *H. asiatica*, and 20 0.3 cm young leaf/normal leaf of *H. asiatica* per DNA extraction). We ground the plant tissues in a 2.0 ml screw tube with one 3.2 mm diameter stainless steel bead and two 2.3 mm diameter zirconia/silica beads (BioSpec Products, Bartlesville, Oklahoma, USA). The tissues were ground at 60 Hz for 30 seconds three times using a STEP tissue grinder (ACTTR, New Taipei City, Taiwan). The DNA sample was extracted following the protocol of the NautiaZ Plant DNA Mini Kit (Nautia gene, Taipei, Taiwan) and eluted with 35 ul UltraPure™ DNase/RNase-Free Distilled Water (Invitrogen, Carlsbad, CA, USA). We used a two-step PCR approach with the ITS3ngs and ITS4 primers following Chen et al. (2021) (Supplementary Table S3) to amplify the nuclear intergenic spacer 2 of the ribosomal DNA (nrITS2). The PCR product was examined on gel electrophoresis and cleaned up by the AMpure XP beads (PCR product: bead 1:1 v:v, Beckman Coulter, Krefeld, Germany). The generated libraries were pooled with equal molar DNA, predetermined by Qubit HS dsDNA HS Assay Kit (Invitrogen, Carlsbad, CA, USA), and sequenced with the Illumina MiSeq platform (300 bp paired-ends) at the NGS High Throughput Genomics Core at Academia Sinica, Taipei. The read data are available in the Short Read Archive (SRA) of NCBI GenBank (BioProject: PRJNA981640).

### Data processing

Raw reads were demultiplexed and the primers were removed by cutadapt [32]. Next, DADA2 [33] was applied to filter out low-quality reads, merge paired-end reads, define unique sequences known as amplicon sequence variants (ASVs), and eliminate chimeric sequences. Subsequently, LULU [34] was utilized for post-clustering of ASVs, removing artificial or intraspecific ASVs that could have arisen from sequencing errors or PCR artifacts. This resulted in a refined ASV table. The taxonomic identity of ASVs was assigned via the RDP Naive Bayesian Classifier algorithm implemented in DADA2, utilizing the UNITE eukaryote database [35]. Finally, the R package “phyloseq” [36] was used to compute alpha diversity indices (observed species, Shannon index) and beta diversity using the Bray-Curtis dissimilarity measure.

### Identification of potential HupA-producing fungi

We determined the potential HupA-producing fungal ASVs with two approaches: 1) sequence clustering, and 2) correlation analyses between HupA content and mycobiome abundance. To gather primary literature on endophytic fungi capable of producing HupA, we conducted a comprehensive literature search in the China National Knowledge Infrastructure (CNKI) database (https://www.cnki.net/) and the Institute of Scientific Information’s Web of Science (WOS) database (https://www.webofscience.com/wos). We focused only on studies that were verified through culture experiments. As of October 20, 2022, we compiled a total of 36 articles reporting HupA-producing fungal strains, comprising 48 strains in total (Supplementary Table S1). We then clustered our ASV sequences with these sequences of known HupA-producing strains by VSEARCH [37] and Geneious Prime 2022.1.1 (https://www.geneious.com) at 97% similarity. As for the correlation analyses, given that the HupA concentration and mycobiomes were not derived from the same plant, we used the mean HupA concentration of three replicates per tissue type to conduct the correlation test (Kendall’s tau rank sum correlation).

### Statistical analysis

To compare the fungal community assemblies in different tissues, we conducted permutational multivariate analysis of variance (PERMANOVA) test with 999 permutations. To test for differences in alpha diversity across tissue types, we conducted ANOVA test with post hoc TukeyHSD test. If the data does not follow a normal distribution, a Kruskal-Wallis test followed by Dunn test was applied instead. The *P* value was corrected by Holm-Bonferroni for multiple comparison corrections. To compare the diversity between the two plant species, a Wilcox rank-sum test was performed. For analysis that required data normalization, we normalized the raw read number to percentage per sample. For alpha diversity measurements, we rarefied the data to 26,328 reads, the lowest number of reads in a sample.

### Nestedness analysis of mycobiome

To investigate mycobiome nestedness, we analyzed the distribution of fungi in different plant tissues of *H. asiatica*. Specifically, the ASVs table was merged by sample tissues, resulting in a presence/absence matrix, in which each row represented a plant part and each column indicated an ASV. A value of “1” indicated the presence of an ASV in a particular plant part, while a “0” indicated its absence. We employed the NODF (Nestedness based on Overlap and Decreasing Fill) metric to quantify the degree of nestedness, ranging from 0 (no nesting) to 100 (perfect nesting) [38]. Additionally, we calculated the nestedness temperature [39], which reflects the degree of nestedness, with lower values indicating a more nested community structure. To validate the accuracy of our nestedness model, we used null models that only preserves the number of presences of ASVs in the matrix while randomizing the distribution across tissues. We performed the data analysis using R packages “bipartite” [40] and “vegan” [41]. To assess the statistical significance of nestedness temperature in our dataset, we conducted 1000 null model simulations.

### Mycobiome comparison between Huperzia asiatica and Diphasiastrum complanatum

To compare the fungal communities between the two species, we combined the two leaf subtypes collected for *H. asiatica* (normal leaf and young leaf) as we did not separate the leaves into subtypes for *D. complanatum*. We then investigated the alpha and beta diversity between the two species as described above. To identify differentially abundant ASVs between *H. asiatica* and *D. complanatum*, we used R packages “DESeq2” [42], which utilizes a negative binomial generalized linear model of reads. As the leaves of *H. asiatica* had a much higher HupA concentration compared to its stems, we focused the differentially abundant test on ASVs of leaves.

### Network Analyses

To further reveal ASVs interactions, networks analyses were carried out. The datasets were split into three categories: A. HupA-rich tissues of *H. asiatica* (including apical shoots, bulbils and young leaves), B. leaves and stems of *H. asiatica*, and C. leaves and stems of *D. complanatum*. ASVs with more than twenty-five sequences of each category were selected to calculate possible correlations. FlashWeave version 0.19.1, a novel co-occurrence method that predicts microbial interaction networks through graphical model inference [43], was used to construct fungal interaction networks, using default parameters. Network properties including R^2^ of power law were calculated in R version 4.2.2. The fast greedy algorithm was used to detect communities [44], based on which, within-module connectivity (Zi) and among-module connectivity (Pi) were calculated to statistically identify microbial keystone ASVs. To assess non-random patterns in the resulting network, we compared our networks against the random networks with equal nodes and edges. Visualization of the constructed networks was performed by Gephi version 0.9.2 [45].

## Results

### Huperzines in tissues of *Huperzia asiatica*

We examined the levels of HupA and its analog HupB, which is considered as a precursor of HupA [46], in different tissues of *H. asiatica* (Fig. 1A). In general, both huperzines showed a similar accumulation pattern in the different tissues of *H. asiatica*: younger tissues had higher huperzine concentrations (Wilcoxon signed rank test: *P* < 0.01; Fig. 1B). For example, the apical shoots of *H. asiatica* had the highest huperzine concentrations (360.27 μg/ml for HupA and 15.00 μg/ml for HupB, respectively), while the normal leaves exhibited the lowest huperzine concentrations (135.90 μg/ml for HupA and 2.00 μg/ml for HupB, respectively). In addition, we found that huperzines were not detected in any tissue of another lycophyte from the Lycopodiaceae, *D. complanatum*, which is consistent with previous reports [16]. These results suggested that *H. asiatica* and *D. complanatum* provided a contrast system to study mycobiome related to HupA biosynthesis.

### Sequencing results

After quality filtering of reads and removal of non-fungal taxa, we obtained 74231 ± 25233 fungal reads for each sample. A total of 925 ASVs were detected (Supplementary Table S4). In *H. asiatica* samples, 441 ASVs were discovered, including 217 in stems, 184 in normal leaves, 183 in young leaves, 97 in sporangia, 56 in bulbils and 55 in apical shoots (Supplementary Fig. S3). In *D. complanatum* samples, 497 ASVs were retrieved, including 319 in leaves and 273 in stems (Supplementary Fig. S3). Only 13 ASVs were commonly shared between the two species.

### Assembly of a list of potential HupA-producing fungi

We further collected the reported HupA-producing fungal strains and their ITS sequences from published literatures. A table containing 48 strains was assembled, most of which belonged to the class Sordariomycetes (Supplementary Table S1). At the genera level, *Colletotrichum* (n = 9), *Fusarium* (n = 6) and *Penicillium* (n = 6) are the most dominant genera. A total of 20 genera, in which these fungi were identified, were subsequently referred as reported HupA-producing (RHP) genera for brevity. By sequence clustering, a total of 24 ASV sequences passing the 97 % sequence similarity cutoff clustered with the reported HupA-producing fungi. Of these 24 ASVs, 18 were identified by both VSEARCH and Geneious, while six were identified only by Geneious (Supplementary Table 5). Furthermore, we performed the correlation analyses between the relative abundance of ASVs and the concentration of HupA across *H. asiatica*’s tissues, by which four ASVs with correlation coefficient (Kendall’s tau b) > 0.4 were identified (Supplementary Table S5). One ASV (ASV10, *Penicillium citrinum*) was identified by both the sequence clustering and correlation analyses. The two analyses rendered a total of 27 ASVs, referred to as potential HupA-producing (PHP) ASVs for subsequent analyses.

### Mycobiome among tissues of *Huperzia asiatica*

We next examined the fungal community in different tissues of *H. asiatica*. Mycobiome structures of *H. asiatica* are significantly different across tissue types (*P* = 0.001; Fig. 2A). The mycobiomes of bulbils and apical shoots are similar to each other while distinct from young leaves, normal leaves, stems, and sporangia (Fig. 2A). Alpha diversity is significantly different across tissue types (*P* < 0.001), with bulbils and apical shoots having the lowest fungal diversity (Fig. 2B). At the fungal class level, bulbils and apical shoots are dominated by Sordariomycetes, while the other tissues have high proportions of Dothideomycetes (Fig. 2C). Based on our PHP list, both Sordariomycetes and Dothideomycetes have been reported to produce HupA. The nestedness temperature of *H. asiatica*’s mycobiomes is 31.478, which is significantly lower than the simulated one (*P* < 0.001), suggesting that the mycobiomes exhibit some degree of nestedness. Stems contain the majority of mycobiomes detected in other tissues (Fig. 2D). Additionally, the matrix fill value is 0.342, which is the mean of non-diagonal elements, indicating that there are relatively few shared ASVs among tissues. The distribution of PHP ASVs (red) is consistent with the overall nestedness structure (Fig. 2D). Stem contains most species of PHP ASVs, while the phyllosphere (apical shoot, bulbil, and leaves) contains some unique PHP ASVs, suggesting that PHP ASVs may originate from different sources.

**Fig. 2.**
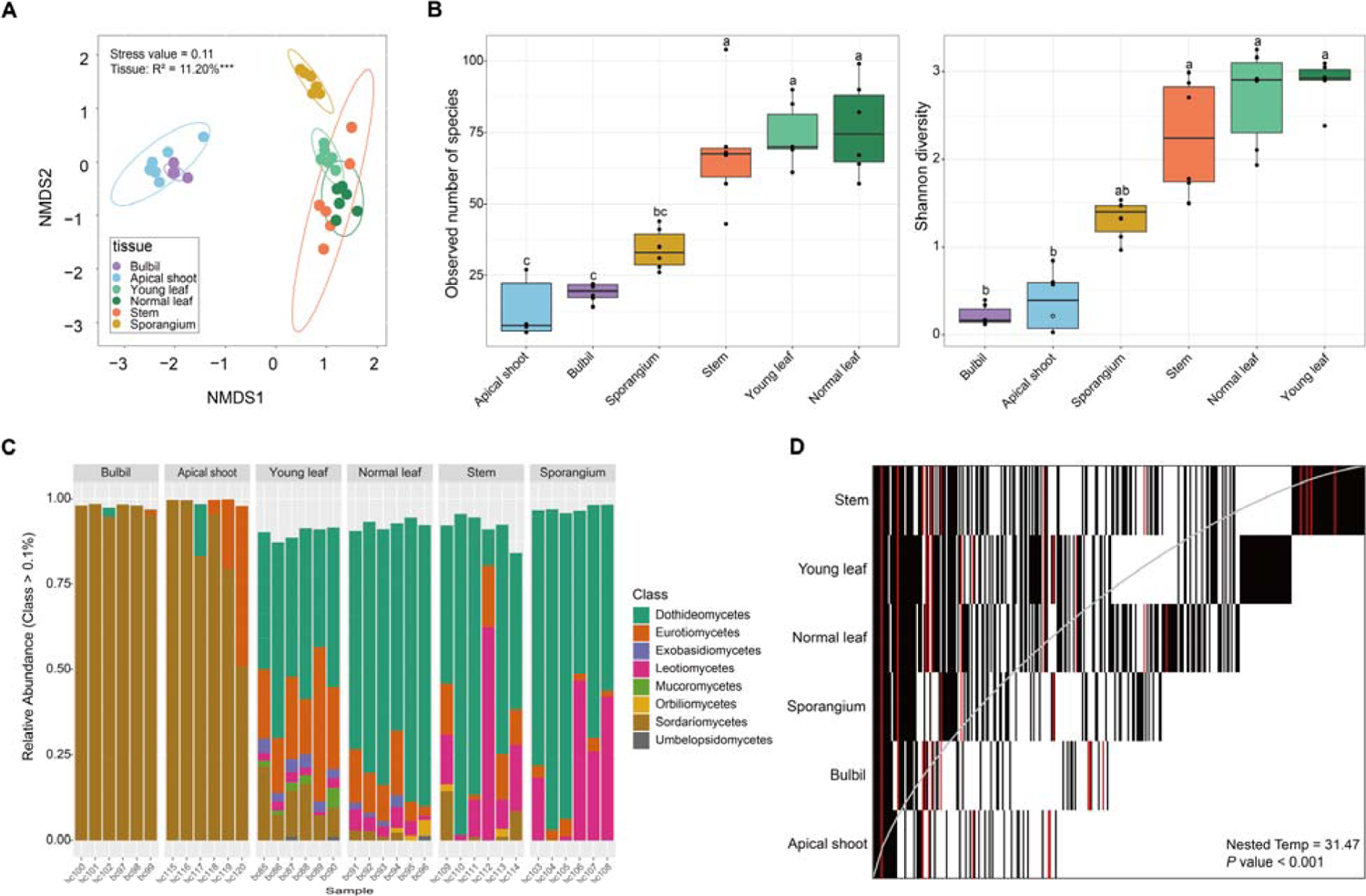
Mycobiome features of *Huperzia asiatica*. **(A)** Non-metric multidimensional scaling (NMDS) ordination plot based on the bray-curtis dissimilarity of *H. asiatica* mycobiomes. *** *P* value = 0.001. Kruskal’s stress values are presented (values less than 0.2 represent good ordination plots). **(B)** Alpha diversity across tissue types in *H. asiatica*. Significant groups in pair-wise comparisons (*p*-adjusted value < 0.05) are indicated by different letters above the boxplots. **(C)** Mycobiome compositions of *H. asiatica* at the class level. The horizontal axis represents the sample name and the vertical axis shows the relative proportion of ASVs annotated to a certain class. The class categories corresponding to each color block are shown in the legend on the right. **(D)** Nestedness plot of fungi aggregated by plant tissues. Each rectangle represents the presence of an ASV in a particular plant tissue. ASVs are arranged from left to right based on their occupancy across plant tissues, and the rows are ordered by decreasing ASV richness from top to bottom. The gray line represents the isocline, and a perfectly nested matrix would have all ASVs on the left side of the isocline.

### Mycobiome comparison between Huperzia asiatica and Diphasiastrum complanatum

The fungal communities of leaf and stem were further compared between the HupA-producing plant *H. asiatica* and the non-HupA producing plant *D. complanatum*. The community assemblies of leaf and stem were significantly different between the two species and two tissue types (*P* < 0.01, Fig. 3A). The number of observed mycobiome species in leaves were significantly higher in *D. complanatum* than that in *H. asiatica* (Kruskal-Wallis test, *P* = 0.02). No differences were detected for Shannon diversity in the two species (Fig. 3B). We then identified the differentially abundant ASVs between the leaves of *H. asiatic* and *D. complanatum*. A total of 71 and 111 ASVs were significantly (*padj* < 0.01, log_2_FoldChange >2) more abundant in *H. asiatica* and *D. complanatum*, respectively. Of the 71 abundant ASVs in *H. asiatica* leaves, seven were identified as PHP ASVs. In contrast, only three out of 111 abundant ASVs in *D. complanatum* were the PHP ASVs. By investigating the genus assignments of the significantly differentially abundant ASVs, six (*Penicillium*, *Trichoderma*, *Alternaria*, *Cladosporium*, *Epicoccum* and *Mucor*, comprising a total of 13 ASVs) and two genera (*Acremonium* and *Leptosphaeria*, comprising a total of 3 ASVs) with reported HupA producing ability were detected in *H. asiatica* and *D. complanatum*, respectively (Fig. 3C).

**Fig. 3.**
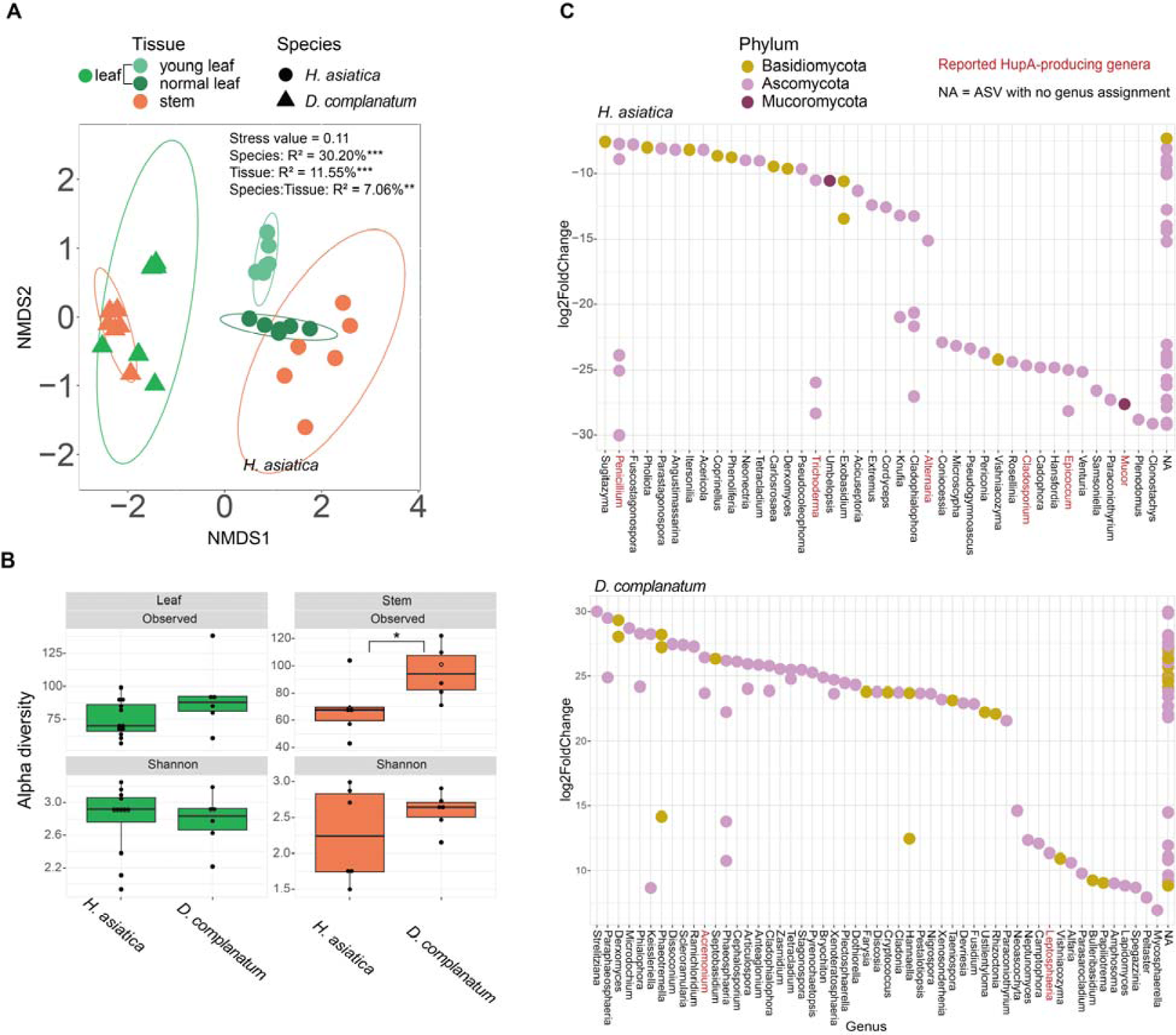
Comparison of mycobiome in the leaves and stems of *Huperzia asiatica* and *Diphasiastrum complanatum*. **(A)** Non-metric multidimensional scaling (NMDS) ordination plot based on the bray-curtis dissimilarity of *H. asiatica* and *D. complanatum* mycobiomes. **(B)** Alpha diversity of endophytic fungi in leaves and stems of *H. asiatica* and *D. complanatum*. **(C)** Differentially abundant taxa between the leaves of *H. asiatica* and *D. complanatum* (Wald test, *padj* < 0.01, log_2_FoldChange > 2). The red font denotes genus with HupA-producing potential in the literature. *** *P* value < 0.001, ** *P* value < 0.01, * *P* value < 0.05.

### Network Analyses revealed potential HupA-producing ASVs’ interactions

We further constructed three interaction networks (A. HupA-rich tissues of *H. asiatica*, B. leaves and stems of *H. asiatica* and C. leaves and stems of *D. complanatum*) to infer their interactions. Supplementary Table S6 showed the main topological properties of the three networks. In all cases, the distribution curves of network node degrees fit well with the power-law model (*R^2^*>0.9, Supplementary Fig. S4). The cohesive characteristics of the network, such as modularity and average clustering coefficient, were higher than those of its corresponding random network, showing the non-random interactions of mycobiome species in the community. These results suggest that these networks conform to the general rule of microbial interaction networks and can be used for subsequent statistical analyses [47–49].

Network A (HupA-rich tissues of *H. asiatica*, i.e., bulbils, apical shoots, and young leaves) consisted of 175 nodes and 166 edges, of which 70.48% were positive edges, indicating a positive correlation between the nodes. We visualized network A based on modules and labeled the PHP ASVs. The network included 28 modules, nine of which contain more than 4% nodes (in detail, 9.14% (module 1), 6.86% (module 4), 6.29% (module 3), 5.71% (module 2), 5.14% (module 5) and 4.57% (module 6, 7, 14, 18) (Supplementary Table S7). Interestingly, some of these PHP ASVs clustered together. For example, ASV1, ASV2, ASV14 and ASV380 were embedded in module 5 and were connected to module 3 containing ASV38 and ASV129 (Fig. 4A). Network B (leaves and stems of *H. asiatica*) consisted of 286 nodes and 274 edges. Network C (leaves and stems of *D. complanatum*) consisted of 363 nodes and 337 edges. Both B and C networks harbor more positive links (90.88% and 91.1%, respectively) than negative links and show high complexity and modularity. However, only five nodes and no edges are shared between the two networks, suggesting that the two plant species have completely different fungi despite that we sampled the same tissue. Consistent with our expectations, *H. asiatica*’s network contained more PHP ASVs than *D. complanatum*’s (13 and 6, respectively).

**Fig. 4.**
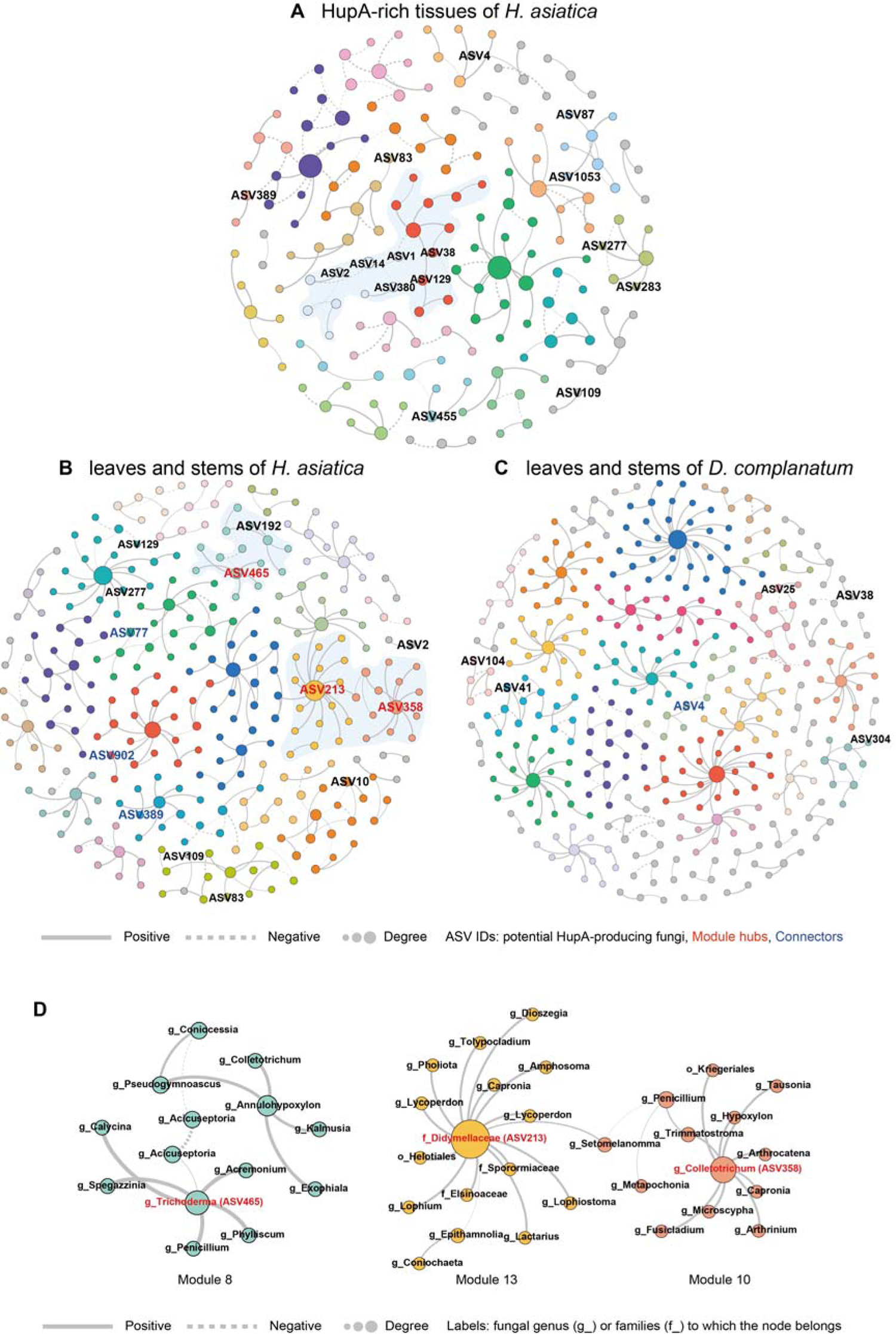
Co-occurrence networks of *Huperzia asiatica* and *Diphasiastrum complanatum* mycobiome community. Networks of **(A)** HupA-rich tissues of *H. asiatica* (including apical shoots, bulbils and young leaves); **(B)** leaves and stems of *H. asiatica*, and **(C)** leaves and stems of *D. complanatum*. The networks are visualized based on modules. Large modules with ≥5 nodes are shown in different colors, and smaller modules are shown in grey. Nodes represent different ASVs and the size of each node is proportional to the number of degrees. Edges refer to the correlation among nodes, being either positive (solid) or negative (dashed). ASV IDs of potential HupA-producing fungi were labeled. **(D)** The modules containing potential HupA-producing fungal ASVs as hubs in the network of leaves and stems in *H. asiatica*. Details of network topological attributes are listed in Supplementary Table S6. The node information for the three networks is in Supplementary Table S7, Supplementary Table S8 and Supplementary Table S9 respectively.

We identified the different roles of the nodes in the three networks based on intra-module connectivity (Zi) and inter-module connectivity (Pi) plots (Supplementary Fig. S5). Most of the ASVs were classified as peripherals (indicating its position in the module in contrast to the hub; 66.86%, 71.33% and 80.99%, respectively), and no network hubs were identified in any of the three networks. *H. asiatica*’s network (network B) has 12 module hubs (including three PHP ASVs) and 70 connectors (including three PHP ASVs) (Supplementary Table S8; Fig. 4B and 4D), while *D. complanatum’s* network (network C) has 10 module hubs and 59 connectors (including one PHP ASVs) (Supplementary Table S9; Fig. 4C). Furthermore, we examined the main fungal genera within modules containing PHP fungi in each of the three networks (Supplementary Fig. S6). The composition of these genera differed between the networks of the two species (*H. asiatica* and *D. complanatum*). Specifically, in both networks A and B of *H. asiatica*, ASVs of the genera *Penicillium* (4 and 5, respectively), *Trichoderma* (4 and 6, respectively), *Dioszegia* (2 and 4, respectively), and *Exobasidium* (2 and 4, respectively) were detected in multiple modules containing PHP fungi. In contrast, in *D. complanatum*’s network (network C), members of the family Didymellaceae (n = 4) and the genera *Phaeosphaeria* (n = 3) exhibited more connections with PHP fungi. These findings highlight the distinct composition of fungal genera associated with PHP fungi between the two species.

## Discussion

The discovery of new fungal species with HupA-producing capabilities involves a series of necessary while laborious processes [28]. Several practical bottlenecks, such as low yield and attenuation of metabolite production during *in vitro* sub-culturing, limit the commercial success of HupA-producing fungi [26]. This study represents the first comprehensive analysis of phyllosphere mycobiomes in *H. asiatica* and *D. complanatum*, two Lycopodiaceae plants with and without capacities to produce HupA, respectively. Moreover, our study offers strategies to increase the likelihood of identifying fungi that could be applied to, or could facilitate, the large-scale commercial production of HupA.

Our study reveals a correlation between HupA content and the diversity distribution patterns of fungi across tissues of *H. asiatica*, consistent with previous studies on *H. serrata* [21]. Notably, we observed a distinct pattern in young tissues, such as bulbils and apical shoots, where the fungal diversity was the lowest despite the highest accumulation of HupA. We postulate that young tissues have limited time for fungal colonization, while increased light exposure due to their positioning on the plant contribute to the high content of HupA [4, 50]. Despite of the low fungal diversity in bulbils, the proportion of potential HupA-producing ASVs is the highest in this tissue (16.07% in bulbils, 10.91% in apical shoot, 5.46% in young leaf, 5.42% in normal leaf, 5.52% in stem, and 5.15% in sporangium). The evidence makes bulbils a promising source for the isolation of HupA-producing fungi, in addition to its function as vegetative propagules [51]. In short, our study raises new candidate tissues for isolating HupA-producing fungi, besides the previously proposed tissues of leaves and stems [20], which encompass the majority of identified HupA-producing strains (Supplementary Table S1) but also comprise other fungal strains enlarging the difficulties of isolation.

The absence of HupA in Lycopodioideae species was observed, but the mycobiome of *D. complanatum* contained several fungi (ASV4, ASV25, ASV38, ASV41 and ASV104) exhibiting high sequence similarity to RHP fungi. These fungi may lack HupA production ability, despite sharing high ITS sequence similarity with known HupA producing strains. The ITS sequences used in the study represent only a small region of the fungal genome, and even minor genetic variations in other parts of the genome could lead to a loss of HupA production capacity [52, 53]. Alternatively, it is also possible that these fungi do possess the ability to produce HupA, but certain specific ecological factors or symbiotic associations necessary for HupA production may be lacking within *D. complanatum* [53]. As a result, the expression of genes related to HupA biosynthesis could be affected. Our network analysis provided further insights into these possibilities by examining the distinct ecological roles played by PHP fungi in the fungal interaction network of both *H. asiatica* and *D. complanatum*. According to Olesen et al. (2007), network hubs (crucial nodes throughout the network with Zi > 2.5 and Pi > 0.62), module hubs (crucial nodes within modules with Zi > 2.5 and Pi < 0.62) and connectors (connecting nodes among modules with Zi < 2.5 and Pi > 0.62) are considered as keystone microbes due to their important connecting roles. These keystone microbes also serve as intermediaries, bridging the host and abiotic factors to plant microbial community [54]. Our results suggest that almost all of the PHP fungal ASVs (4 out of 5) were peripherals (Non-critical nodes with Zi < 2.5 and Pi < 0.62) in the network of *D. complanatum*. In contrast, almost half of the PHP ASVs (6 out of 13) occupied key positions in the network of *H. asiatica* as module hubs and connectors (Fig. 5). Through complex interactions among microorganisms, they may play a vital role in system-level coordination for HupA production and overall fungal community dynamics. Furthermore, the notable interactions between PHP fungi and other fungi within the network or the same module indicated new clues for searching potential fungal facilitators. The common method to find potential fungal facilitators is to combine fungi that have similar ecological niches, since they colonize in close physical distance and embedded in one complex microbial community [30]. Therefore, fungal ASVs in the same module as PHP fungal ASV may serve as fungal facilitators, mitigating the decline in HupA-producing capacity through co-culture. By examining the distribution of these fungal genera across different networks, we can make a more informed assessment of their potential as facilitators. For instance, ASVs of the genera *Penicillium* and *Trichoderma* were consistently detected in more than four modules containing PHP fungi in both *H. asiatica* networks (Supplementary Fig. S6), making them the most promising candidates for future co-cultivation investigations. Additionally, ASVs of the genera *Dioszegia*, *Exobasidium*, *Lycoperdon* and *Cladosporium* were also recommended as they were detected in four modules containing PHP fungi in at least one of the *H. asiatica* networks. On the contrary, caution should be exercised with fungal genera or families such as *Phaeosphaeria* and Didymellaceae, as they were predominantly associated with modules containing PHP fungi in the *D. complanatum* network, suggesting they may not be suitable facilitators.

**Fig. 5.**
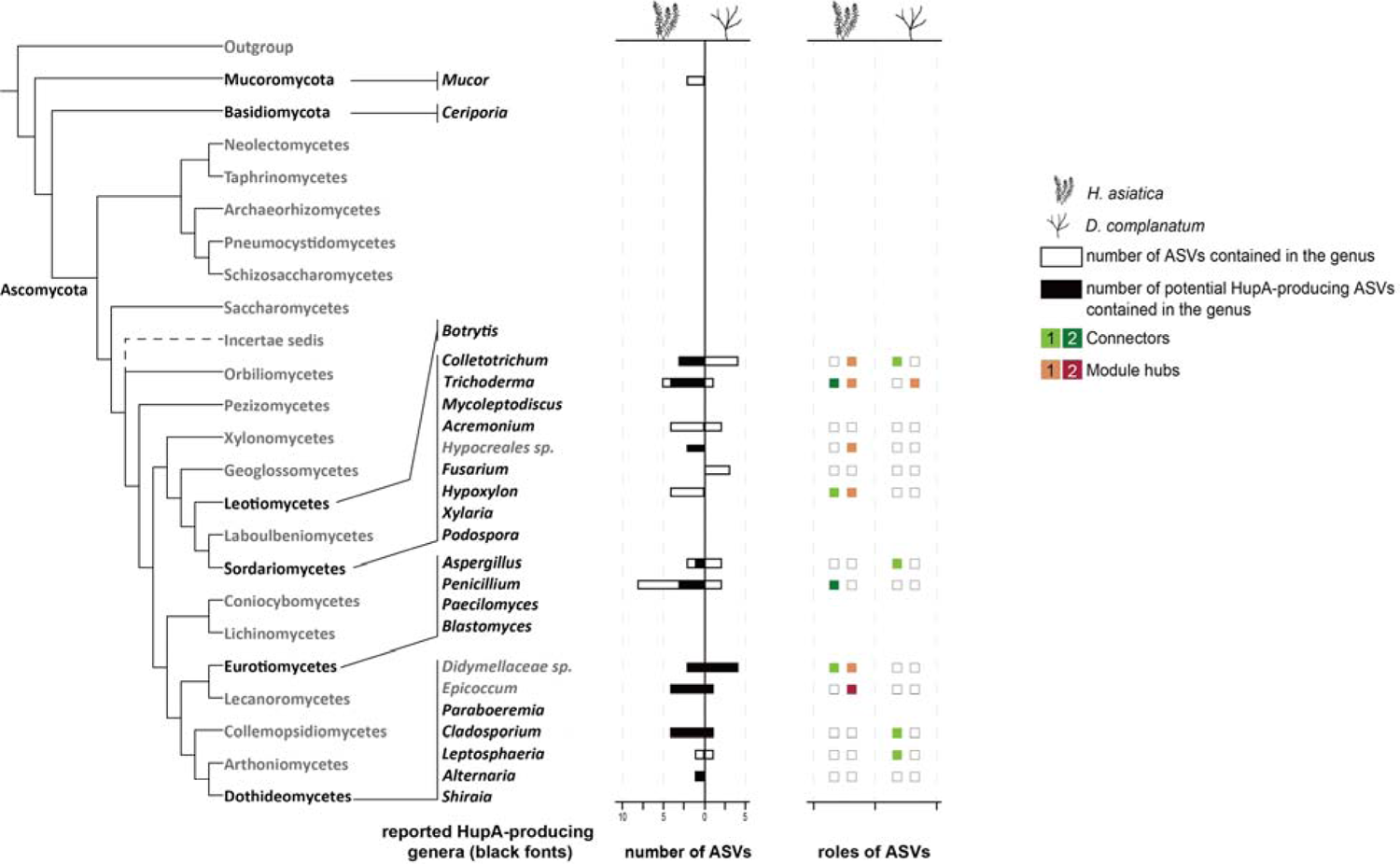
The roles of potential Huperzine A producing fungi in mycobiome networks. Phylogenetic distribution of the reported HupA-producing genera (black fonts). Empty horizontal bars imply the number of ASVs contained in the genus, and the horizontal bars filled with colours indicate the number of potential HupA-producing ASVs. The colors of the squares show the role of HupA-producing ASVs in the network, with green indicating the connectors, red indicating the module hubs, and shades of colour indicating the number of ASVs in each category.

It is important to acknowledge that our experimental pipeline did not exclude epiphytic fungi, and thus the observed fungal taxa in our sequencing efforts includes both endophytic and epiphytic fungi, which can reside on the plant surface as well as within plant tissues [55]. Previous studies have highlighted distinct differences between phylloplane (leaf surface) and endophytic fungal communities within the same leaves [56]. Epiphytic fungi have been reported with specific secondary metabolic potential, such as taxol producing potency [57]. It’s possible that some epiphytes are responsible for HupA production. Therefore, further detailed analyses are necessary to differentiate endophytes from epiphytes and explore their potential contributions to HupA production.

## Conclusions

This study advances our knowledge of phyllosphere fungal diversity in Lycopodiaceae species and provides valuable insights into the search for potential HupA-producing fungi and fungal facilitators. It highlights the notable variations of HupA concentration and mycobiome assemblies across tissues, pressing the importance of exploring young tissues. Furthermore, we revealed that the potential HupA-producing fungi are often hubs in the mycobiome networks of HupA-producing plants, implying that the ecological interactions among microbes may play roles in mitigating and sustaining the fungi-mediated production of complex bioactive compounds. Future investigations into these intricate interactions will deepen our understanding of the molecular and ecological factors influencing HupA biosynthesis and its industrial application.

## Declarations

### Ethics approval and consent to participate

The acquisition of plant materials in this study adheres to pertinent institutional, national, and international guidelines and regulations. Plant sample collection was approved by Changbai Mountain Ecological Protection Station (*H. asiatica*) and Forestry and Nature Conservation Agency I-lan Branch (*D. complanatum*). The voucher specimen of *H. asiatica* was deposited in SZG (Herbarium, Fairy Lake Botanical Garden, Shenzhen & Chinese Academy of Sciences, No. 00120204), and *D. complanatum* was deposited at TAIF (Herbarium of Taiwan Forestry Research Institute, No. 545100).

### Consent for publication

Not applicable.

### Availability of data and materials

The short read data are available in the Short Read Archive (SRA) of NCBI GenBank (BioProject: PRJNA981640). Code for sequence processing and statistical analysis is presented at GitHub (https://github.com/koshroom/Huperzia_mycobiome).

## Authors’ contributions

K.C. and L.W. conceived and designed the study. L.K, K.C., and L.W. provided the plant materials. R.D. measured the huperzine concentrations. K.C. and C.C. carried out the amplicon sequencing. C.L. performed metabolome data analysis. K.C. analyzed the amplicon data. L.L. conducted the network analysis. L.L., J.J., and K.C. visualized the data. L.L. wrote the first draft of the manuscript with specific contributions from K.C., L.W., C.L., and R.D., and all authors read and approved the final draft.

## Acknowledgements

We thank Dr. Yao-Moan Huang for fieldwork assistance and Dr. Fay-Wei Li for valuable discussion regarding this project.

## Supplemental Material

Supplementary materials are posted as submitted.

Supplementary Tables (S1-S9): HupA_Supplementary Tables.xlsx

Supplementary Figures (S1-S6): HupA _Supplementary Figures.docx

## Supplementary Figures for

**Figure S1.**
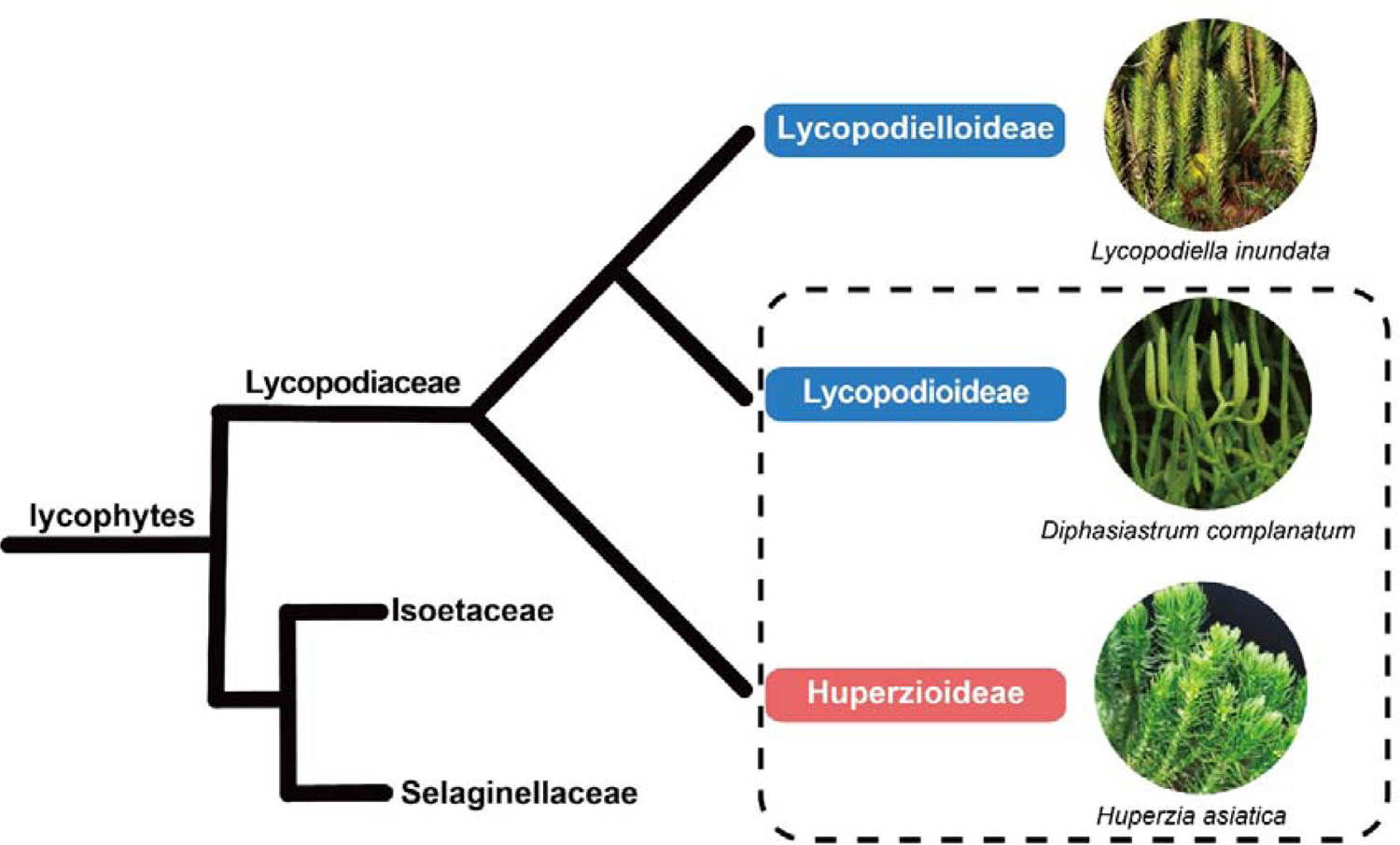
Phylogeny of lycophytes. The blue-shaded subgroups do not produce HupA, and the red-shaded one can produce HupA. The dashed lines frame the materials involved in our study. Image of *D. complanatum* courtesy of P.-F. Lu. Image of *L. inundata* courtesy of Wikipedia.

**Figure S2.**
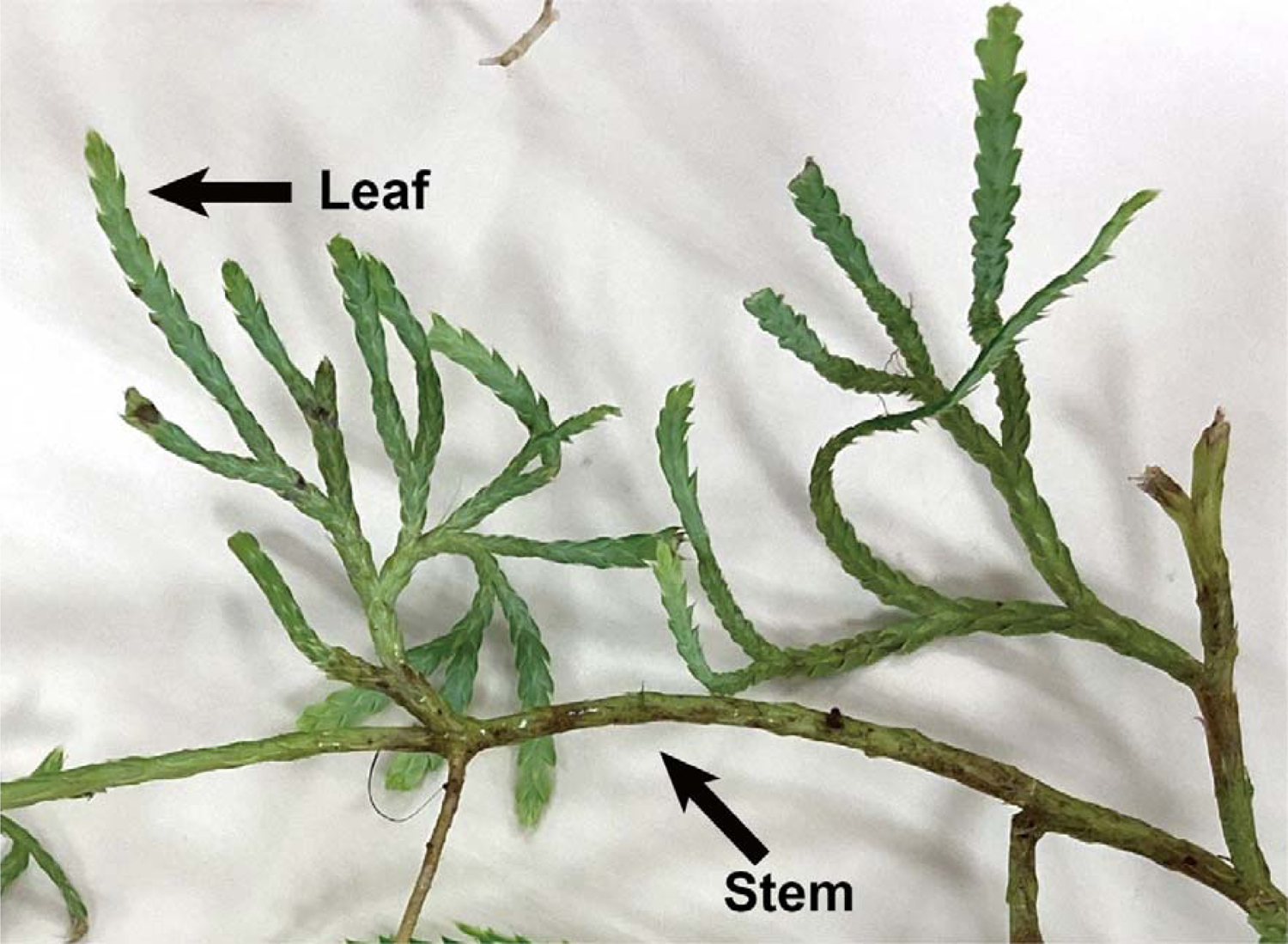
Stem and leaf of *Diphasiastrum complanatum*

**Figure S3.**
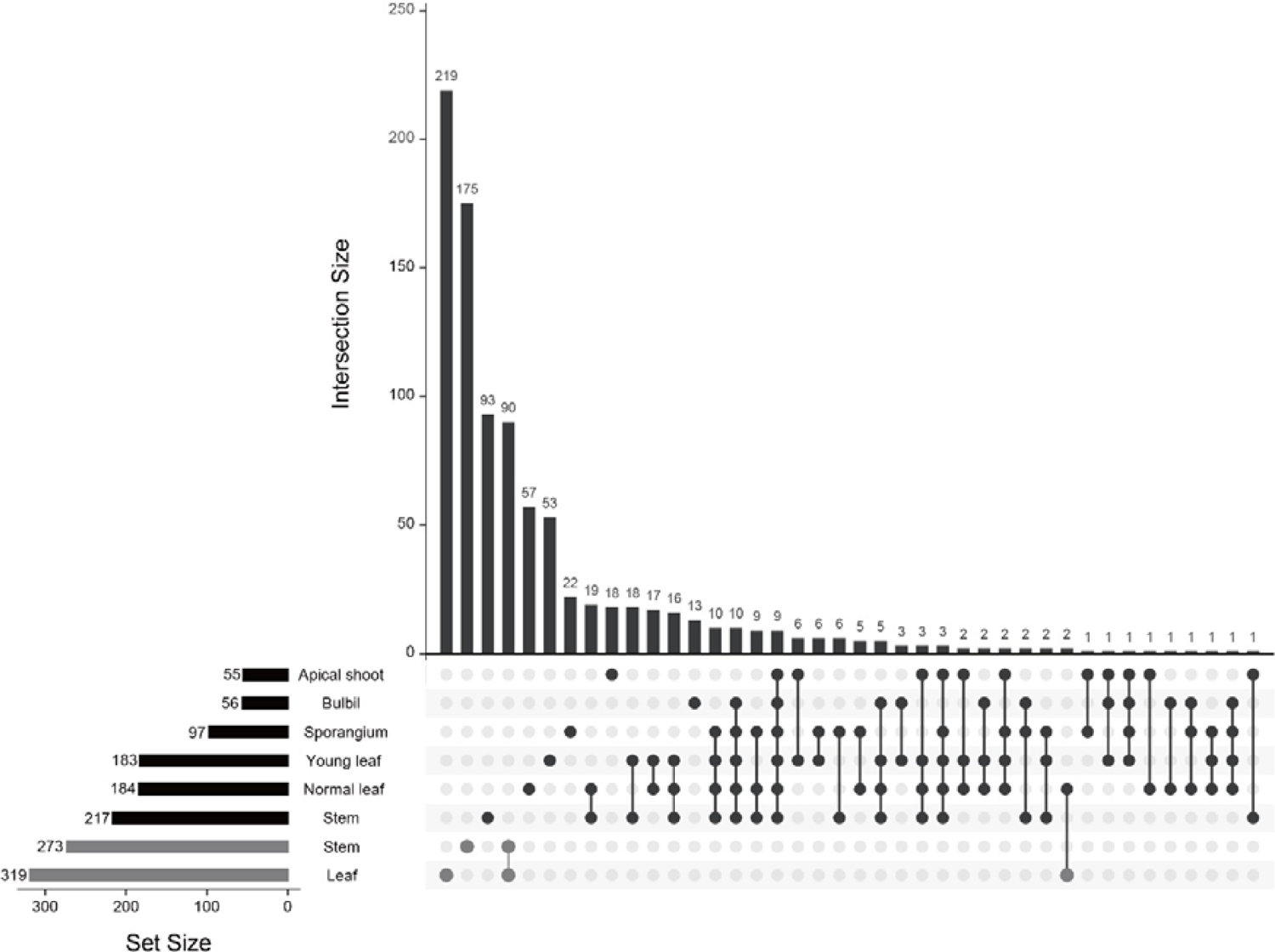
UpSet plot showing the unique and intersected ASVs among samples. Total ASVs in each sample are displayed through set size, in which black bars are sets of *Huperzia asiatica* and gray bars are sets of *Diphasiastrum complanatum*. The vertical bars represent the distinct and overlapping ASVs in different tissues, and dots below indicate the samples included in each vertical bar.

**Figure S4.**
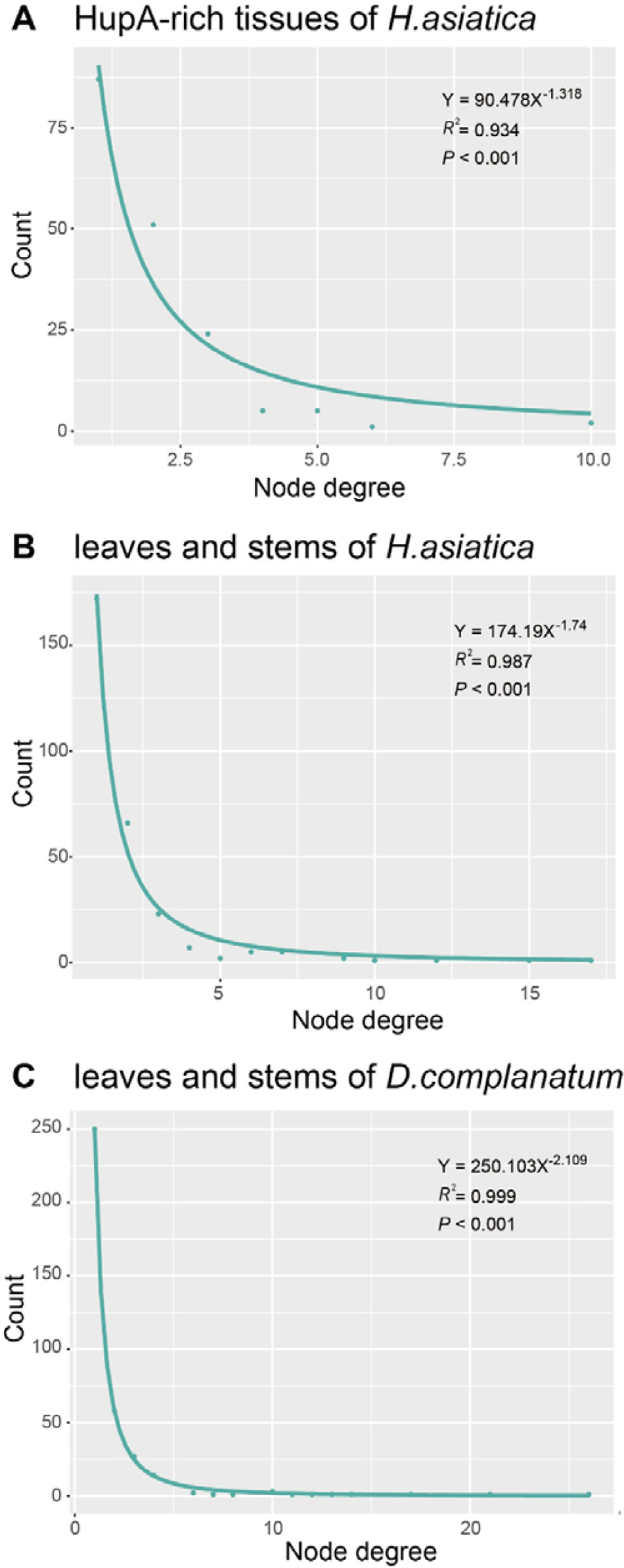
The distribution curves of network node degrees fit with the power-law model.

**Figure S5.**
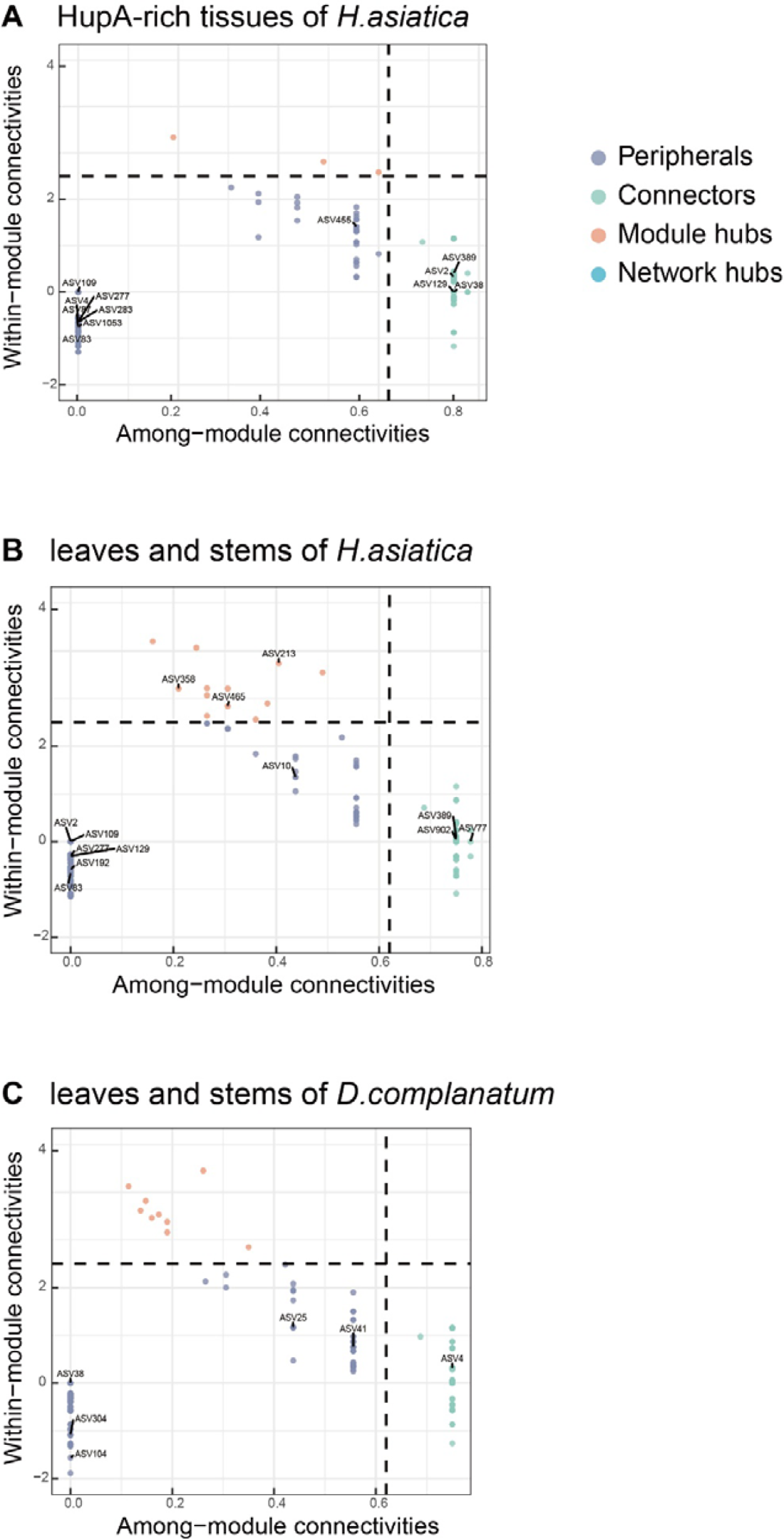
Intra-module connectivity (Zi) and inter-module connectivity (Pi) plots.

**Figure S6.**
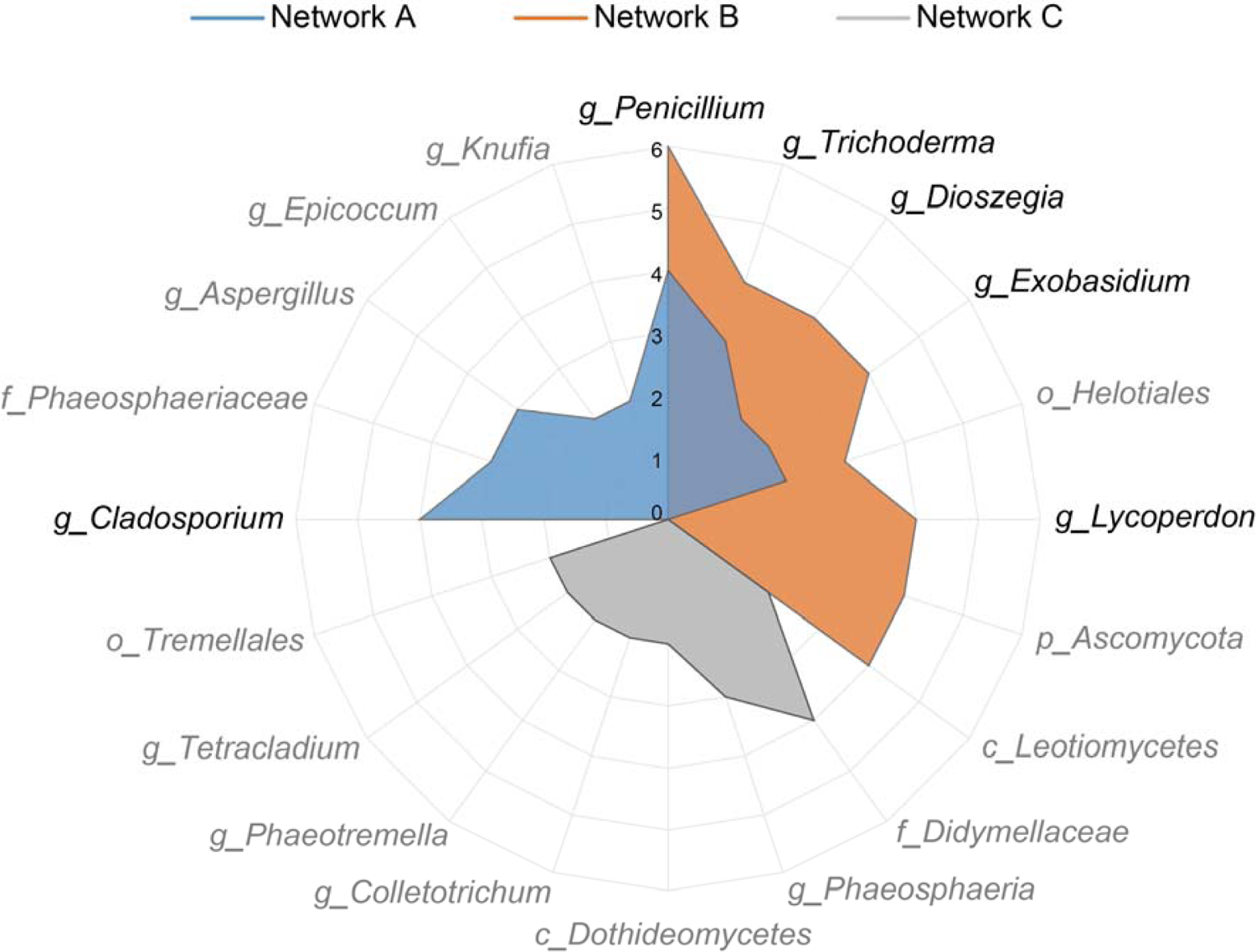
A radar plot comparing the number of ASVs belong to the main fungal genera within modules containing PHP fungi in network A (HupA-rich tissues of *Huperzia asiatica*, i.e., bulbils, apical shoots, and young leaves), network B (leaves and stems of *H. asiatica*) and Network C (leaves and stems of *Diphasiastrum complanatum*). The black font labels candidates that we believe are promising for future co-cultivation investigations (i.e. was detected in at least four modules in Network A or B).

